# Modelling Sex Differences in Circadian Regulation of Kidney Function of the Mouse

**DOI:** 10.1101/2022.08.26.505440

**Authors:** Anita T. Layton, Michelle L. Gumz

## Abstract

Kidney function is regulated by the circadian clock. Not only do glomerular filtration rate (GFR) and urinary excretion oscillate during the day, the expressions of several renal transporter proteins also exhibit circadian rhythms. Interestingly, the circadian regulation of these transporters may be sexually dimorphic. Thus, the goal of this study is to investigate the mechanisms by which kidney function of the mouse is modulated by sex and time of day. To accomplish this, we have developed the first computational models of epithelial water and solute transport along the mouse nephrons that represent the effects of sex and circadian clock on renal hemodynamics and transporter activity. We conduct simulations to study how the circadian control of renal transport genes affects overall kidney function, and how that process differs between male and female mice. Simulation results predict that tubular transport differs substantially among segments, with relative variations in water and Na^+^ reabsorption along the proximal tubules and thick ascending limb tracking that of GFR. In contrast, relative variations in distal segment transport are much larger, with Na^+^ reabsorption almost doubling during the active phase. Oscillations in Na^+^ transport drive K^+^ transport variations in the opposite direction. Model simulations of BMAL1 knockout mice predict a significant reduction in net Na^+^ reabsorption along the distal segments in both sexes, but more so in males than females. This can be attributed to the reduction of mean ENaC activity in males only, a sex-specific effect that may lead to a reduction in blood pressure in males.

## Introduction

Most physiological systems exhibit circadian rhythms that are synchronized with solar time and organize with a 24-hour period. In mammals, the circadian rhythms are mediated in part by the central clock, which resides in the suprachiasmatic nucleus (SCN) of the hypothalamus and acts as a central pacemaker for rhythms observed in behavior, body temperature, and hormones (1). Another key player in the circadian system are peripheral clocks that are present in most organs (e.g., liver and kidney) and cells that can oscillate independently of the SCN (2). The central and peripheral clocks essentially share the same molecular architecture (3,4). The core clock genes Bmal1 and Clock dimerize to induce the transcription of target genes, including the Period (Per) and Cryptochrome (Cry) genes. PER and CRY then heterodimerize to act on the CLOCK-BMAL1 protein complex, thereby inhibiting their own transcription and completing the main feedback loop (5). In terms of kidney function, disruption of the circadian clock in animal models results in changes in blood pressure, alterations in the circadian pattern of water and electrolyte urinary excretion, and altered sensitivity to dietary salt challenges (6,7).

In humans and animals, glomerular filtration rate (GFR), filtered electrolyte loads, urine volume, and urinary excretion exhibit significant diurnal rhythms (8–10). Those rhythms reflect the regulation of kidney function by the circadian clock; specifically, clock proteins BMAL1 and PER1 regulate a number of transporter proteins in the kidney, including the Na^+^/H^+^ Exchanger 3 (NHE3), Sodium-Glucose co-Transporter 1 (SGLT1), Epithelial Na^+^ Channels (ENaC), pendrin, and Renal Outer-Medullary K^+^ channels (ROMK) (11–13), which likely contributes to the circadian regulation of renal function. The expressions of some of these transporters follow a pattern similar to GFR and peak during the active phase (e.g., pendrin and ENaC), whereas others peak during the rest phase (e.g., NHE3 and ROMK) (11).

Interestingly, Crislip et al. demonstrated that the regulation of renal Na^+^ transport by BMAL1 differs between male and female mice (11). Indeed, findings of this study provide further evidence that physiological studies must take into account two variables: time of day and sex. Notably, sex has impact on physiological functions that are regulated by, and those that are independent of, the circadian clock. The kidney mass and single-nephron GFR (SNGFR) of a male mouse are both larger than female (14,15). Perhaps more remarkably, Veiras et al. reported that the renal electrolyte transporters and channels that mediate electrolyte transport in rodent kidneys have abundance patterns that are sexually dimorphic (16). Notably, female mice exhibit a lower NHE3 abundance in the proximal tubules, and thus lower transport capacity, compared to males. The higher fractional Na^+^ distal delivery in females is handled by the augmented transport capacity in the downstream segments, including a higher abundance of Na^+^-K^+^-Cl^-^ co-transporter (NKCC), Na^+^-Cl^-^ co-transporter (NCC), and claudin-7 (16). As already noted, the expressions of many of these transporters are regulated by the circadian clock, in a sexually dimorphic manner (11).

### How does the circadian control of renal transport genes affect overall kidney function, and how does that process differ between male and female mice?

To answer these questions, we simulate circadian rhythms in key renal transport activities using sex-specific computational models of water and electrolyte transport along the nephrons of the male and female mouse kidneys. Prior modeling efforts have focused on humans and rats (not mice), to study the alternations in renal transport processes under various dietary conditions (17–19), pathophysiological conditions and therapeutic manipulations (20–22), and pregnancy (23). Recently, sex-specific models were developed to capture the sexual dimorphism in renal hemodynamics and transporter expression patterns in rats (24,25) and humans (26,27). However, almost all published models are based on the steady-state formulation and do not consider circadian rhythms in transporter activities and kidney function. A notable exception is Ref. (22) which couples a computational model of the renal circadian clock to the activity of the NHE3 in an epithelial transport model of the proximal convoluted tubule cell of the rat kidney. A limitation of that study is that it represents a stand-alone proximal convoluted cell, and does not consider the circadian oscillations in other transporters, nor can it predict urinary excretions. Also, the model is formulated for the male rat, which is not directly applicable if one wishes to understand findings in knockout studies in male and female mice (11). Hence, in this study we develop comprehensive sex-specific computational models of circadian regulation of epithelial transport of solute and water transport in the mouse kidney. Using those models, we conduct simulations to answer the following questions: *How does the differential circadian regulation of the expression levels of key transporter genes impact the transport processes along different nephron segments during the day? And how do those effects differ between males and females?*

## Methodology

We conduct simulations using our recently published sex-specific models of the mouse kidney (28). A schematic diagram of the various cell types is given in Fig. 1. These models represent the sexual dimorphism in size, renal hemodynamics, and transporter expression patterns in mice (16,29) and the circadian regulation of GFR and transporter activities(8). We simulated the protocol with mice kept in a light-dark cycle. GFR and selected transporter activities are set to vary as sinusoidal functions of time. Model parameters that exhibit circadian rhythms are summarized in Table 1 and Fig. 2.

**Table 1.**
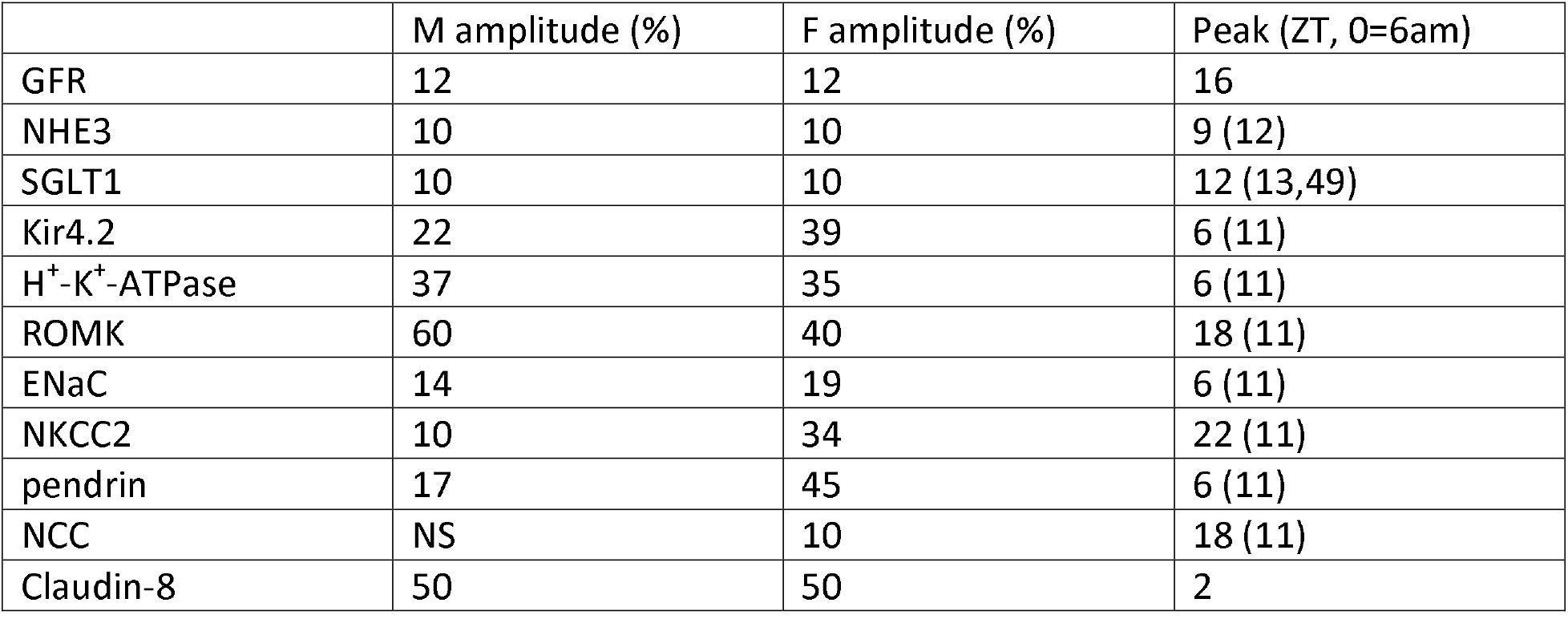
Peak times and oscillation amplitudes of model parameters that exhibit circadian rhythms.

|  | M amplitude (%) | F amplitude (%) | Peak (ZT, 0=6am) |
| --- | --- | --- | --- |
| GFR | 12 | 12 | 16 |
| NHE3 | 10 | 10 | 9 (12) |
| SGLT1 | 10 | 10 | 12 (13,49) |
| Kir4.2 | 22 | 39 | 6 (11) |
| H<sup>+</sup>-K<sup>+</sup>-ATPase | 37 | 35 | 6 (11) |
| ROMK | 60 | 40 | 18 (11) |
| ENaC | 14 | 19 | 6 (11) |
| NKCC2 | 10 | 34 | 22 (11) |
| pendrin | 17 | 45 | 6 (11) |
| NCC | NS | 10 | 18 (11) |
| Claudin-8 | 50 | 50 | 2 |

**Figure 1.**
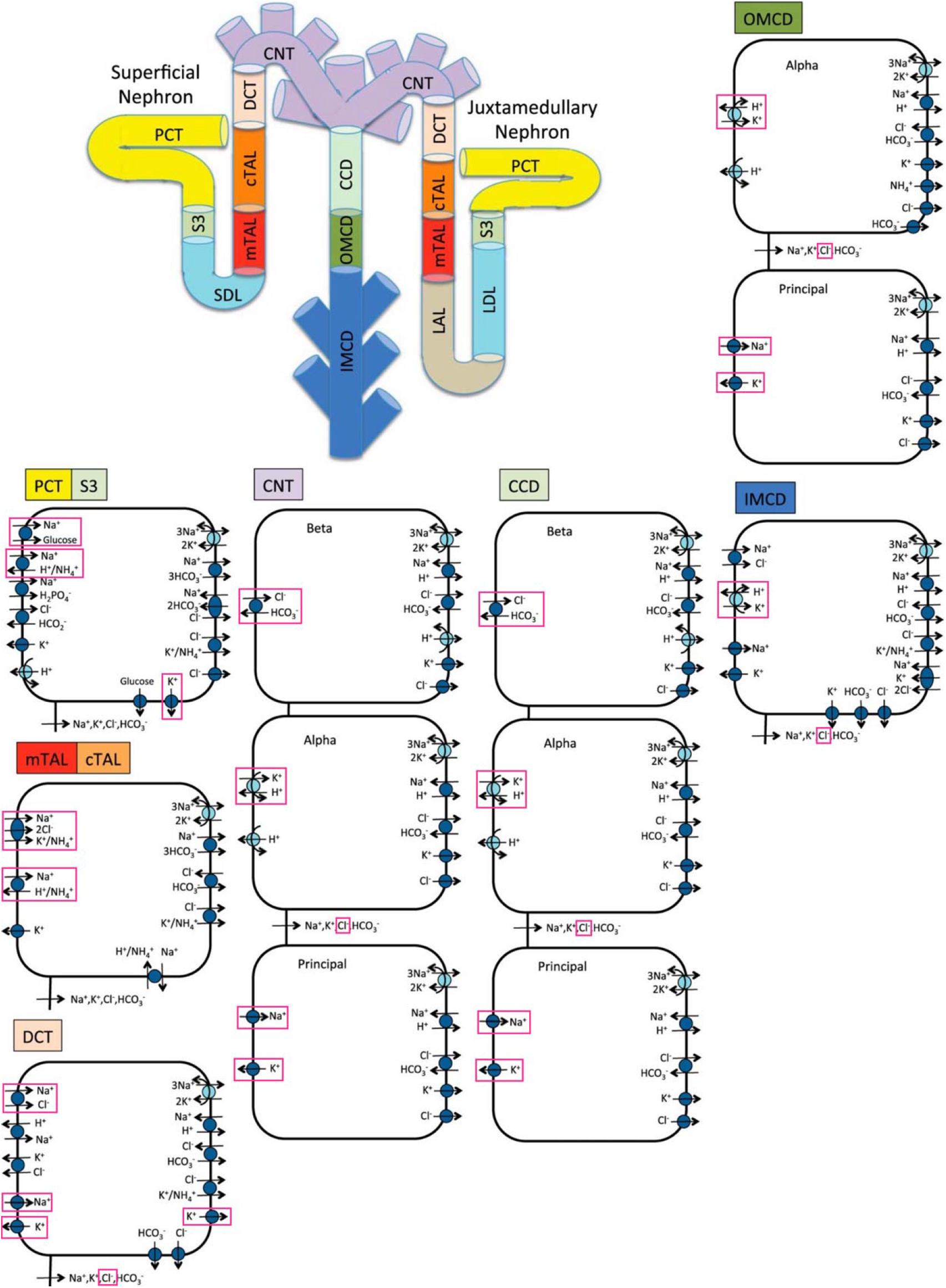
Schematic diagram of the nephron system (not to scale). Model structure is similar for the rat and mouse. The model includes one representative superficial nephron and five representative juxtamedullary nephrons, each scaled by the appropriate population ratio. Only the superficial nephron and one juxtamedullary nephron are shown. Along each nephron, the model accounts for the transport of water and 15 solutes (see text). The diagram displays only the main Na^+^, K^+^, and Cl^-^ transporters. Transporters and channels that are assumed regulated by the circadian clock are highlighted in red. PCT, proximal convoluted tubule; SDL, short or outer medullary descending limb; mTAL, medullary thick ascending limb; cTAL, cortical thick ascending limb; DCT, distal convoluted tubule; CNT, connecting tubule; CCD, cortical collecting duct; OMCD, outer-medullary collecting duct; IMCD, inner medullary collecting duct; LDL, thin descending limb; LAL, thin ascending limb. Adopted from Ref. (50).

**Figure 2.**
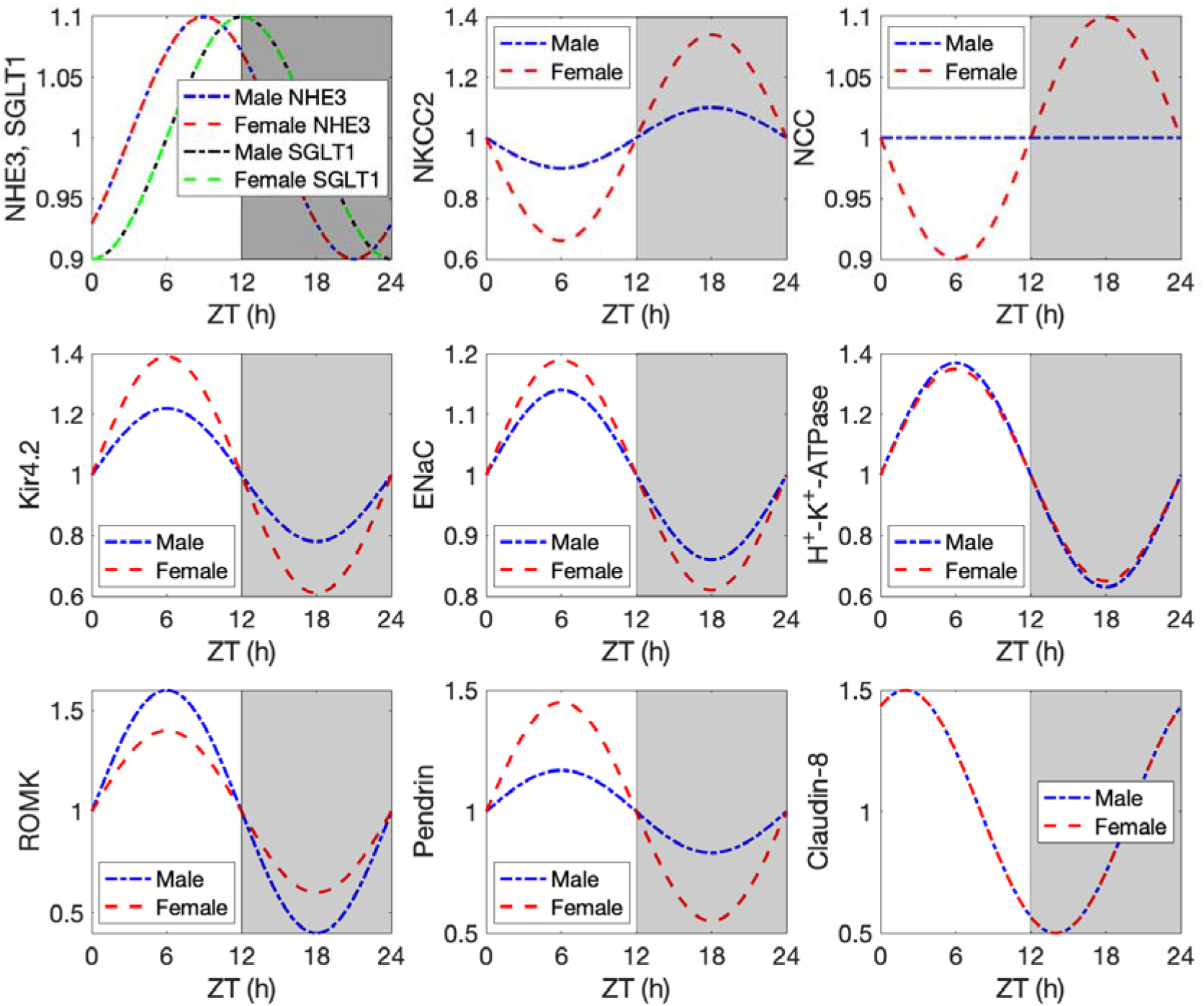
Time-profiles of circadian regulated transporter activities. Normalized to mean values.

Not only are renal electrolyte transporters and channels that mediate electrolyte transport regulated by the renal circadian clock, they also exhibit abundance patterns that are sexually dimorphic (16,30). As previously noted, female mouse proximal tubules exhibit a lower NHE3 abundance and activity. Consequently, relative to males, in females the proximal tubule reabsorbs a substantially lower fraction of filtered Na^+^ (16). The higher fractional Na^+^ distal delivery in females is handled by the augmented transport capacity in the downstream distal tubular segments. Female mice exhibit higher abundance of NKCC, NCC, and claudin-7 along the distal nephron segments (16).

### Model structure

Mouse kidneys consist of primarily superficial and juxtamedullary nephrons, with a ratio estimated to be 82:11 in male mice (31). In the absence of data, we assume the same ratio in females. The loops of Henle of juxtamedullary nephrons reach into differing depths of the inner medulla. To capture this heterogeneity, we represent six classes of nephrons: a superficial nephron (denoted by “SF”) and five juxtamedullary nephrons that are assumed to reach depths of 1, 2, 3, 4, and 5 mm (denoted by “JM-1”, “JM-2”, “JM-3”, “JM-4”, and “JM-5”, respectively) into the inner medulla. The ratios for the six nephron classes are taken to be *n*_*SF*_ = 0.82, *n*_*JM*−1_ = 0.82 × 0.4 *n*_*JM* − 2_ = 0.82 × 0.3, *n*_*JM* − 3_ = 0.82 × 0.15, *n*_*JM* − 4_ = 0.82 × 0.1, and n_*JM* − 5_ = 0.82 × 0.05 (31). The model superficial nephron includes the proximal tubule, short descending limb, thick ascending limb, distal convoluted tubule, and connecting tubule segments. Each of the model juxtamedullary nephron includes all the same segments of the superficial nephron with the addition of the long descending limbs and ascending thin limbs; these are the segments of the loops of Henle that extend into the inner medulla. The length of the long descending limbs and ascending limbs are determined by which type of juxtamedullary nephron is being modeled. The connecting tubules of the five juxtamedullary nephrons and the superficial nephron coalesce into the cortical collecting duct. SNGFR for juxtamedullary nephrons is assumed to be 20% higher than the superficial nephron SNGFR, based on superficial SNGFR and GFR measurements (15,32).

Each nephron segment is modeled as a tubule lined by a layer of epithelial cells in which the apical and basolateral transporters vary depending on the cell type (i.e., segment, which part of a segment, intercalated and principal cells). The models account for the following 15 solutes: Na^+^, K^+^, Cl^-^, HCO_3_^-^, H_2_CO_3_, CO_2_, NH_3_, NH_4_^+^, HPO_4_^2-^, H_2_PO_4_^-^, H^+^, HCO_2_^-^, H2CO_2_, urea, and glucose. The models consist of a large system of coupled ordinary differential and algebraic equations, solved for steady state, and predicts luminal fluid flow, hydrostatic pressure, membrane potential, luminal and cytosolic solute concentrations, and transcellular and paracellular fluxes through transporters and channels.

### Bmal1 knockout simulations

We simulated the sex-specific alterations in the diurnal changes in distal transporter and channel expression reported in Ref. (11). Specifically, mean ENaC activity was reduced by 26% in males only. Mean pendrin activity is reduced by 40% and 10% in males and females, respectively; additionally, oscillation amplitude in males changes from 17% to - 20%. Mean ROMK expression increases by 40% and 24% in males and females, respectively.

## Results

We conducted model simulations to predict solute and water transport along nephrons in the male and female mouse kidneys, at a range of zeitgeber times (ZT). GFR and filtered Na^+^ and K^+^ loads are shown in Figs. 3A1-3A3. Using these as inputs, the models predicted fluid and solute flows along all nephron segments. The flows of fluid, Na^+^, and K^+^ into key nephron segments (proximal tubules, loops of Henle, and distal convoluted tubules) are shown at selected ZT in Fig. 4. Predicted water transport, Na^+^ transport, and K^+^ transport along individual nephron segments are shown in Fig. 5, where the “distal tubule” segment includes the distal convoluted tubule, connecting tubule, and collecting duct.

**Figure 3.**
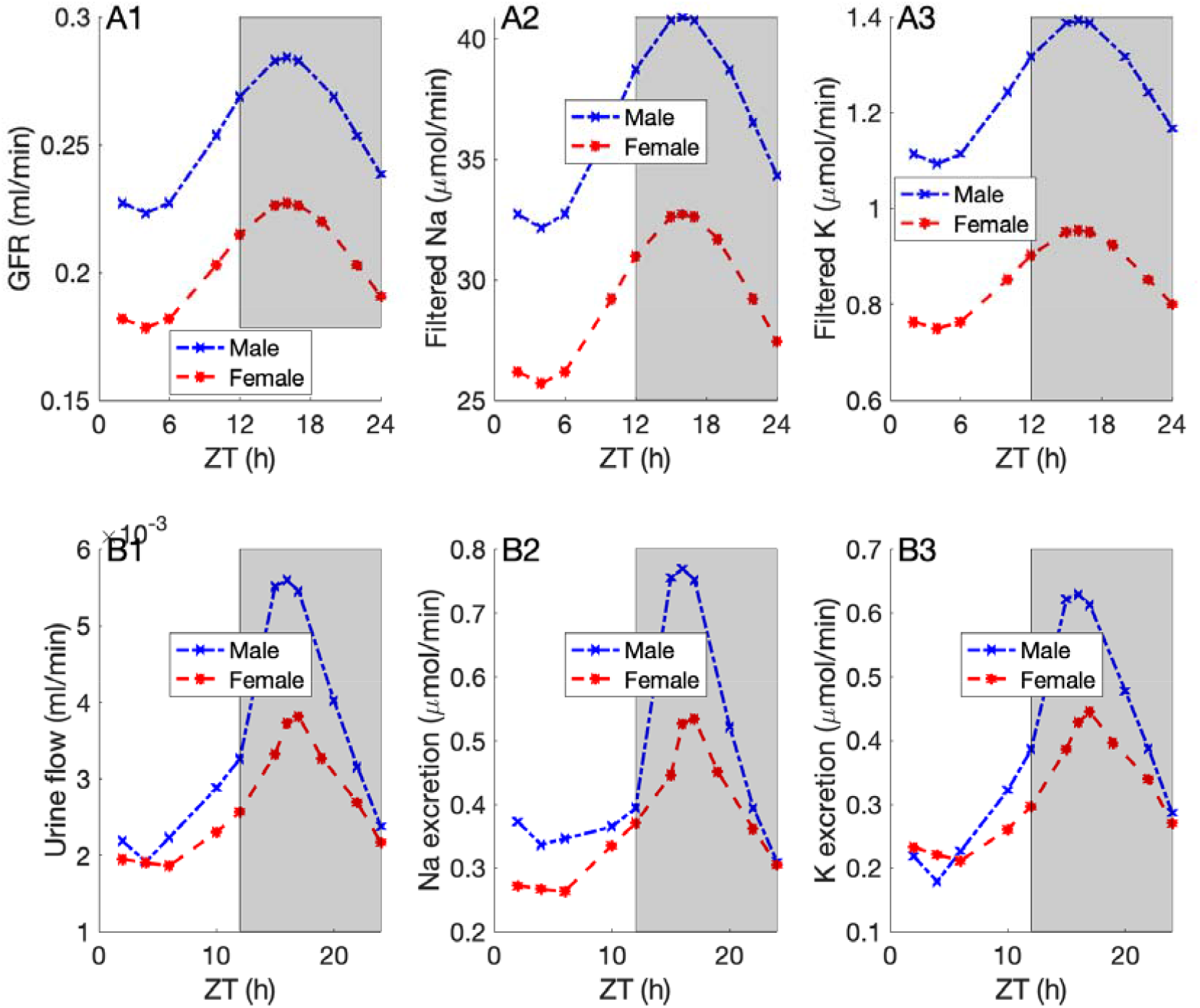
Prescribed time profiles of GFR and filtered Na^+^ and K^+^ loads (A1 – A3). Predicted urine output, Na^+^ excretion, and K^+^ excretion (B1 – B3).

**Figure 4.**
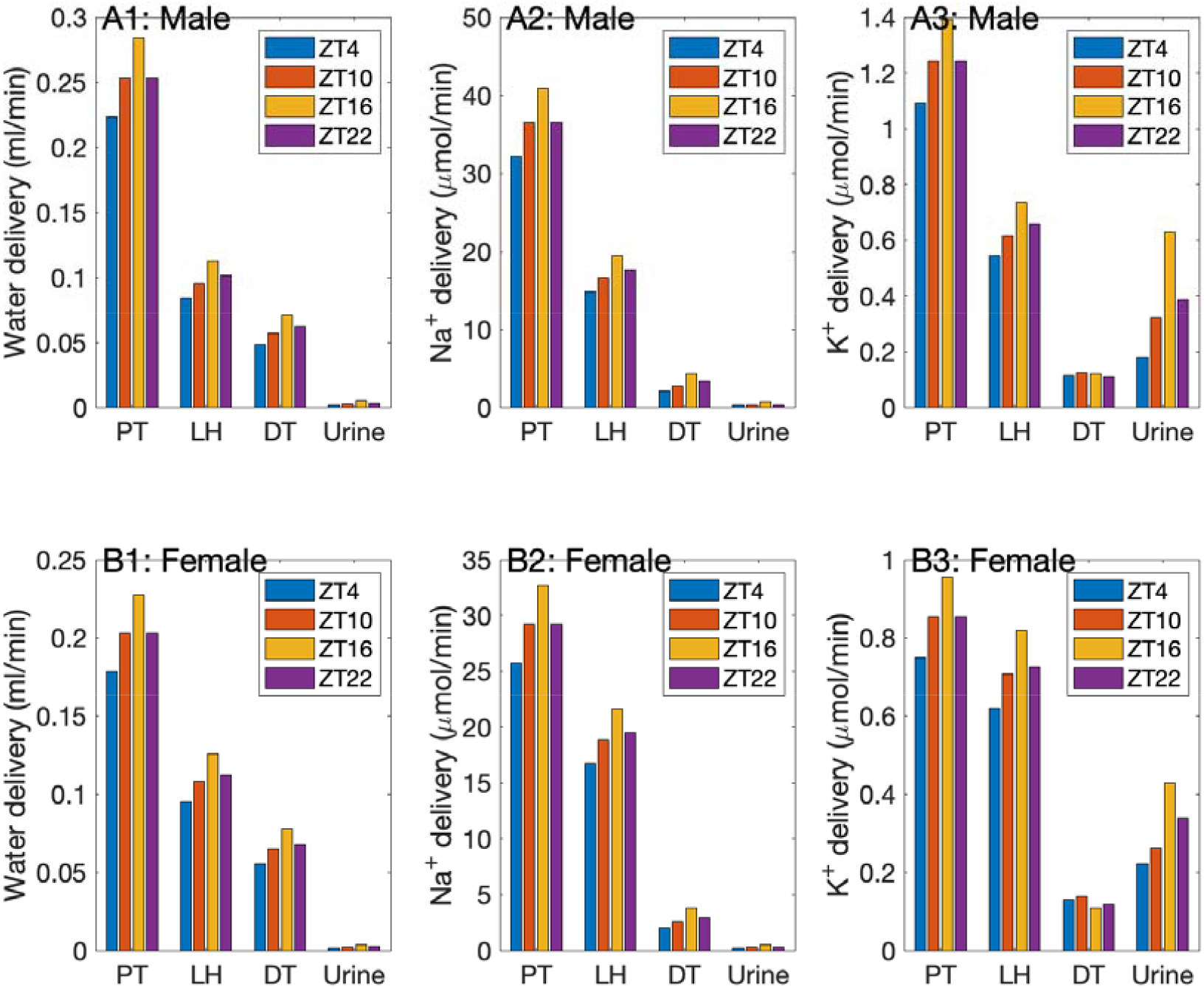
Predicted segmental deliveries, obtained for male (A1-A3) and female (B1-B3), at different zeitgeber times. Shown are flows of water, Na^+^, and K^+^ into the proximal tubules (PT), loops of Henle (LH), and distal convoluted tubules (DT), and also urine excretion rates.

**Figure 5.**
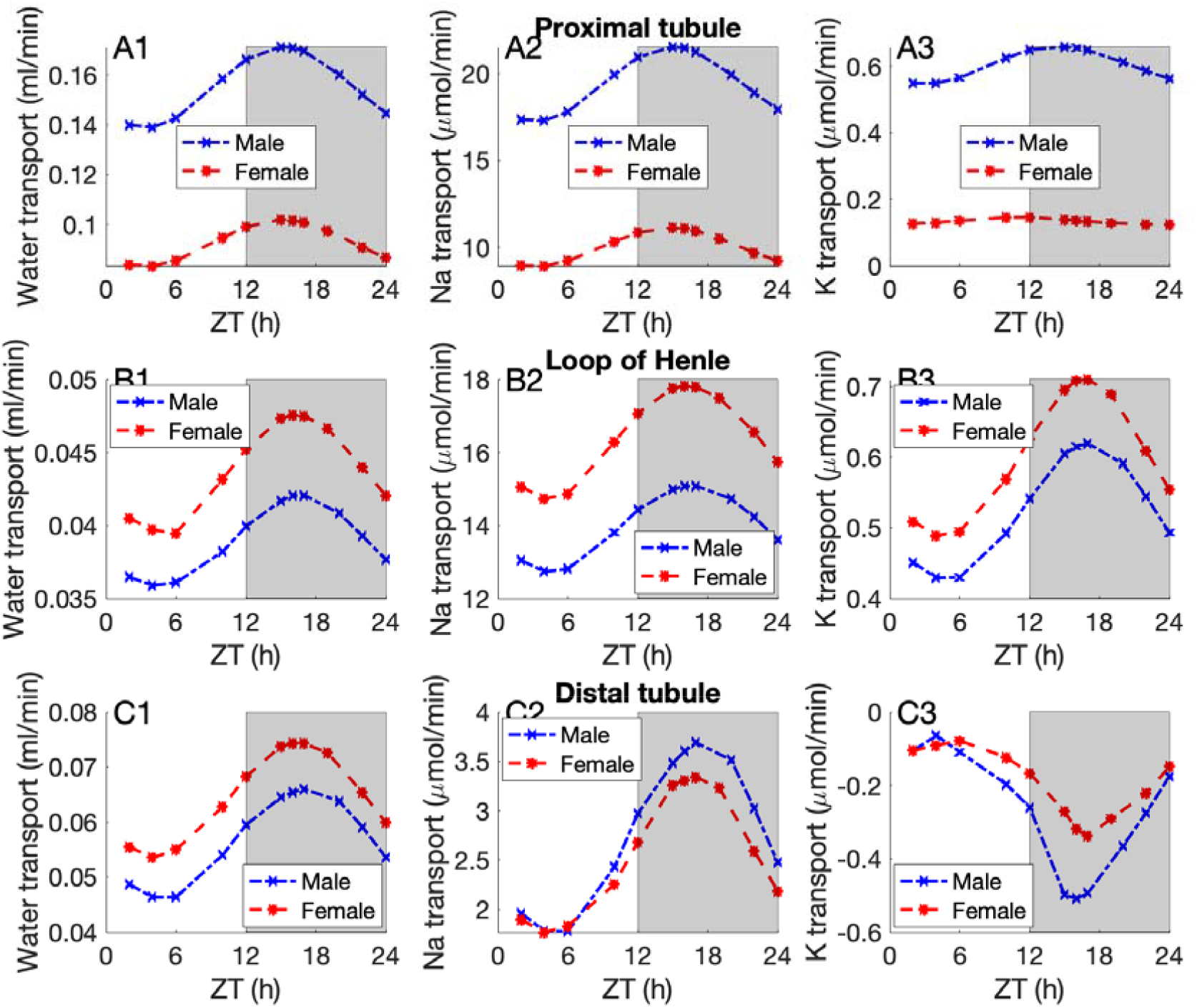
Predicted segmental transport, obtained for male (blue) and female (red), at different zeitgeber times. Transport taken positive for reabsorption.

We first summarize predicted segmental flows and transport in broad strokes. The male mouse model predicts that about half of the filtered Na^+^ and K^+^ is reabsorbed along the proximal tubules; electrolyte transport drives water reabsorption. Most of the remaining electrolytes are reabsorbed along the thick ascending limbs. The distal segments reabsorb most of the remaining Na^+^ and water, but secrete K^+^. Throughout the day, segmental transport changes as filtered loads change, so that the fractional segmental transport remains relatively constant. A qualitatively similar pattern is predicted by the female mouse model, except that due in large part to their lower NHE3 activity, the proximal tubules in the female model transport significantly less Na^+^, K^+^, and water, with the downstream thick ascending limbs and to a lesser extent, distal nephron segments, picking up the load. Taken together, the models predict urine output and excretions that are similar between the two sexes during the inactive phase, but substantially higher in the male during the active phase (Figs. 3B1-3B3).

### Renal circadian clock differentially regulates segmental transport in a sex-dependent manner. How do the circadian rhythms in renal hemodynamics and transporter activities translate into variations in tubular transport and eventually urine excretion?

Circadian rhythms in GFR and transporter activities are assumed to have the same phase relations in both sexes; however, the oscillation amplitudes may be different between males and females. See Table 1. Recall that GFR is assumed to exhibit diurnal variations of 12%, with a peak at ZT16 (Table 1, Fig. 3A1). Also modulating proximal tubule Na^+^ and water transport are oscillations in NHE3 and SGLT1, which are out of phase with SNGFR, peaking at ZT9 and ZT12, respectively (Table 1). The models predict that in both male and female the circadian oscillations in proximal tubule transport are almost in-phase with SNGFR but with a slightly larger amplitude of 12.5% due to the contributions of NHE3 and SGLT1; see Figs. 5A1-5A3. Because the proximal tubules in the male mouse kidney mediate a larger fraction and net amount of the filtered volume and solutes, the net oscillation amplitude is significantly larger in males compared to females.

Along the thick ascending limbs, NKCC2 exhibits circadian oscillations almost in-phase with GFR, with a significantly larger amplitude in female (Fig. 2). Consequently, while the Na^+^ transport along that segment varies in both sexes, the diurnal oscillation amplitude is significantly larger in females (Figs. 5B1-5B3). Along the distal tubular segments, several transporters exhibit circadian rhythms, with most peaking during the inactive phase at ZT6. But the effect of filtered load variations dominates, and the circadian oscillations in distal tubule Na^+^ and water transport are close to GFR in phase (Figs. 5C1-5C2). K^+^ transport is driven by Na^+^ transport and oscillates in the opposite direction (Fig. 5C3). Because the distal tubular segments transport only a small fraction of the filtered loads, the relative amplitude of these oscillations are much larger than GFR. The amplitude of the distal Na^+^ transport oscillations is 54 and 44%, respectively, in the male and female models (Fig. 5C2). Even more drastic changes can be seen in distal K^+^ secretion, which varies by a factor of 4 in female, and a factor of 8 in males (Fig. 5C3). These diurnal variations in segmental transport result in urine volume and excretions are that 2-3 times higher during the active (dark) phase (Figs. 3B1-3B3), consistent with reported values (33,34).

### Bmal1 in the distal segments regulates Na^+^ handling in a sex-specific manner

Crislip et al. reported in kidney-specific cadherin BMAL1 knockout mice sex-specific changes in Na^+^ and K^+^ transporter expressions along the distal nephron segments. We simulated solute and water transport along the nephrons of male and female BMAL1 knockout mice; see Methods. The models predict a reduction in net Na^+^ reabsorption along the distal segments (distal convoluted tubules to collecting ducts) in both sexes, but more so in males (0.73% of filtered load) than females (0.44%). This is due in large part to the 26% reduction in mean ENaC activity, assumed in males only (11). While the changes in distal Na^+^ transport are rather miniscule, note that urinary Na^+^ excretion is only ∼15% of total distal Na^+^ transport. As such, the models predict a 4.2% increase in urinary excretion in males, but only 0.5% in female. While neither natriuretic response is drastic, it is likely enough at least in the male to cause a reduction in blood pressure (11).

### Circadian rhythms in urine output and salt excretions are driven primarily by GFR

The circadian clock mediates a host of coordinated changes in renal hemodynamics and transporter expressions. To assess the extent to which these two types of changes contribute to the circadian variations in urinary excretions, we conducted simulations in which changes in GFR and transporter activities are selectively eliminated. Model predictions are summarized in Fig. 6 for three cases: circadian rhythms intact in both GFR and transporter expressions (labelled ‘Control’), transporter activities only (‘Transport’) with GFR set to their steady-state values (28), and GFR only (‘GFR’) with transporter activities set to their mean values.

**Figure 6.**
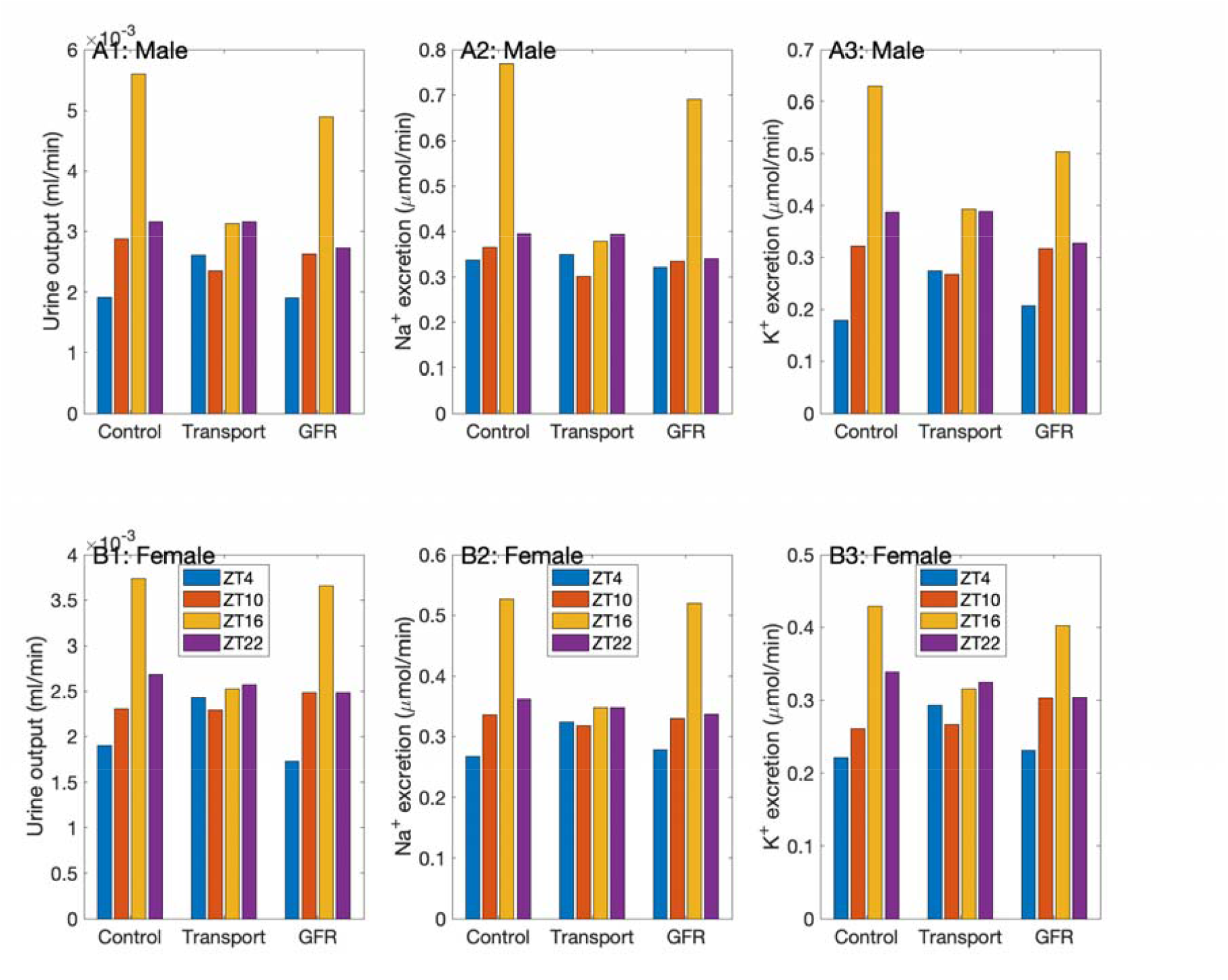
Effect of circadian rhythms in renal hemodynamics and transporter expression on kidney function. Comparison of urine output and excretions with circadian rhythms intact in both GFR and transporter expressions (‘Control’), transporter expressions only (‘Transport’), and GFR only (‘GFR’). Results obtained for male (A1-A3) and female (B1-B3), at selected zeitgeber times.

A comparison of predicted urine output and excretions indicates that for both male and female, changes in urine output and excretions are driven primarily by circadian oscillations in GFR, especially at high filtration rates (ZT16). If GFR remains constant throughout the day, the circadian rhythms in transporter expression would yield variations in urinary excretions that are out of phase with control, with a peak closer to ZT20. That is a result of the competing effects of transporter activities that peak during the day (NHE3, Kir4.2, ENaC, H^+^-K^+^-ATPase, ROMK, pendrin, claudin-8) versus those that peak during the night (NKCC2, NCC). Taken in isolation, the peaks of the activities of some major transporter (e.g., NHE3, ENaC), which mediate Na^+^ reabsorption, are expected to correspond to the troughs of urine output and Na^+^ excretion. Thus, as the activities of NHE3 and ENaC peak during the day, urinary excretions peak at night in the ‘Transport’ case.

## Discussion

Essentially every aspect of biological function changes according to time of day. In the context of kidney function, blood pressure, filtration rate, transporter expression, and urinary output have all been reported to exhibit circadian rhythms. Nevertheless, variations across the day are often ignored in the design and reporting of research. In recent years, there has been an increasingly number of experimental studies focusing on the circadian oscillations or diurnal changes in kidney function (e.g., Refs. (6,7)). In contrast, computational models of kidney function have largely ignored the effect of diurnal variations. This is the first study that incorporated not only time of day as a variable, but sex as well, into computational models of the mouse kidney.

The present models represent the 24% oscillations in GFR, as well as the sex-specific variations in transporter expression levels (11). The models predict that urine output and excretions change by a factor of 2-3 (Fig. 3), consistent with observations (6,10). Recall that urine output and Na^+^ excretion account for about 1% of filtered values. Thus, despite the large fluctuations in urinary excretions, circadian oscillations in overall transport of water and electrolytes parallel GFR and vary by 20%. That is accomplished by tubuloglomerular balance, whereby the proximal tubules absorb approximately the same fraction of the filtered loads, as well as variations in key transporters.

By means of model simulations, we seek to gain insights into processes that are difficult to study *in vivo*. Specifically, we determine the variations in water and electrolyte transport along individual nephron segments. Simulation results suggest that such variations differ substantially among segments and between sexes. In both males and females, variations in water and Na^+^ reabsorption along the proximal tubules track that of GFR, with a predicted peak-to-trough difference of approximately 25%. This tubuloglomerular balance is driven in part by the flow-dependent transport of the proximal tubules. Similarly, thick ascending limb transport also track oscillations in GFR, due to variations in Na^+^ delivery (Figs. 4A2-4B2) and NKCC2 activity (Fig. 2). In contrast, variations in distal segment transport are much larger, with Na^+^ reabsorption almost doubling during the active phase. That is a result of the elevated Na^+^ delivery at night (Figs. 4A2-4B2), an effect that dominates the concurrent circadian suppression of ENaC (Fig. 2).

Quantitative immunoblotting studies have indicated that NKCC2 and NCC are more abundant in female versus male mice (16). Interestingly, the amplitudes of the circadian oscillations of these two transporters are substantially larger in females (Table 1). Hence, when taken in isolation, sex differences in nephron transport that are mediated by the differential expressions of NKCC2 and NCC may be augmented during the night and attenuated during the day. Indeed, model simulations predict that the difference in Na^+^ reabsorption is augmented during the active phase along the thick ascending limbs (Fig. 3B2) and the distal convoluted tubules (results not shown). It is noteworthy that while distal Na^+^ transport is higher during the active phase, ENaC activity is elevated during the rest phase. This may contribute to the higher efficacy of therapies that manipulate ENaC, such as ACE inhibitors, when taken in the evening (rest phase for humans) (35). Natriuretic and diuretic responses following the administration of benzamil were found to be significantly greater in male compared with female rats whether given at the beginning of the rest phase or active phase (36). A better understanding of the circadian regulation of kidney function, and the underlying molecular mechanisms, might allow us to identify the optimal time to administer certain medications.

Kidney-specific knockout studies have reported that loss of renal BMAL1 lowers blood pressure in male mice but not in female mice (11,37). To understand the mechanism that explains the differential impact of BMAL1 knockout, we simulated water and electrolyte transport in the BMAL1 male and female mouse kidneys. These simulations represent the reported effect of BMAL1 knockout on renal transporter activities; in particular, ENaC activity is significantly suppressed in males but not in females (11). Consequently, the model predicts negligible change in urinary Na^+^ excretion in female, but a small increase (4%) in males. That minor but significant natriuretic response in the male mouse may be sufficient to explain the drop in blood pressure (11). This sex difference can be attributed to the attenuated ENaC activity that is, as noted above, found in male BMAL1 knockout mice only. If ENaC activity were not reduced in the male BMAL1 knockout model, but the changes in pendrin and ROMK were kept intact, the natriuretic response in males would be essentially eliminated (simulation results not shown).

Results of the present study join an expanding body of experimental works to highlight the important roles of sex and time of day in regulating physiological function. Thus, variations between sex and across the day be taken into account not only in the design and reporting of biomedical research, but in modelling analysis as well. One particularly interesting application is hypertension. There are well-known sex differences in blood pressure regulation, which have implications on blood pressure, on the prevalence of hypertension, and on the pathogenesis and progression of hypertension-related cardiovascular and renal disease (38–40). Moreover, blood pressure follows a distinct circadian pattern, characterized by a marked decline during sleep (usually approximately 10–20% lower than the mean daytime value) followed by a surge in the early morning (41). That morning rise in arterial blood pressure coincides with the peak incidence of many cardiovascular events (42). To unravel the complex interactions between the effects of sex and time of day on blood pressure regulation and related diseases, one may utilize computational modeling techniques. The present models can form the basis of integrative sex-specific blood pressure regulation models that represent the effects of circadian rhythms. To that end, the present models must be extended. While these models are highly detailed, one major limitation is that the interstitial fluid compositions are assumed known a priori; as such, the interactions among nephron segments and vasculature are not represented. The vasculature is particularly relevant in this context, in part because of known sex differences in the bioavailability of nitric oxide, a powerful vasodilator, in spontaneously hypertensive rats (43). Thus, the present models can be incorporated into a full kidney model that captures the interactions among the model nephrons by determining the interstitial fluid concentrations based on solute and water reabsorption from tubular segments (44–46). The resulting comprehensive sex- and time-of-day-specific kidney function models can then be incorporated into a sex-specific blood pressure regulation model (47,48) expanded to represent circadian rhythms.

